# The effects of chronic neuropathic pain states on the discriminative stimulus effect of fentanyl and other MOR agonists

**DOI:** 10.1101/2024.11.19.624329

**Authors:** Gwendolyn E. Burgess, John R. Traynor, Emily M. Jutkiewicz

**Affiliations:** Department of Pharmacology, University of Michigan, Ann Arbor Michigan, 48109

## Abstract

Pleasant subjective effects of drugs (e.g., euphoria) have been demonstrated to contribute to their abuse potential. In humans, there is some evidence that acute pain states may decrease the positive subjective effects of opioids; however, no studies have directly tested the impact of a long-lasting pain state. Therefore, the goal of this study was to directly evaluate the discriminative stimulus of mu opioid receptor (MOR) agonist, fentanyl, or the non-opioid drug of abuse, cocaine, in the presence or absence of spared nerve injury (SNI) induced chronic neuropathic pain. Prior to surgery, MOR agonists (fentanyl, morphine, nalbuphine) dose-dependently increased % fentanyl-like responding, as expected; surprisingly, after surgery, we saw small, significant rightward shifts in the fentanyl and morphine dose response curves in *both* sham and SNI groups suggesting that the observed shifts were not due to chronic pain. In both sham and SNI groups, there was an increase in the generalization of nalbuphine to the fentanyl-discriminative stimulus. There was no change in the discriminative stimulus of cocaine (or amphetamine substitutions) over 4 months of SNI-induced chronic neuropathic pain or sham states, suggesting that the SNI model failed to alter the discriminative stimuli of fentanyl and cocaine. Following induction of chronic neuropathic pain, there was an observed increase in quinpirole-induced generalization to the cocaine discriminative stimulus. In the future, studies should directly examine the abuse potential of low efficacy MOR agonists and dopaminergic agonists in the presence and absence of chronic pain states.

**Significance Statement:** Subjective or interoceptive effects of drugs of abuse are known to contribute to the abuse potential. This study demonstrated that long-lasting neuropathic pain failed to alter the discriminative stimulus of mu opioid receptor agonists or cocaine; however, we observed an increase in quinpirole-induced generalization to the cocaine discriminative stimulus, suggesting the abuse potential of direct dopaminergic agonists should be further evaluated in the presence or absence of pain states.

## Introduction

Opioid analgesics are known to produce both pleasant (e.g., “euphoric”), and unpleasant (e.g., “dizzy”, “nauseous”) subjective effects in humans (Comer et al., 2008; 2009; 2010; Lasagna et al., 1955; Zacny, 2001). In non-human animals (non-verbal), a correlate of subjective effects is measured through use of drug discrimination assays, and there is generally a similarity in observed drug effects across species (Schuester & Johansen, 1976; Porter et al. 2018). While the internal state produced by drug may not be evident to others, these private, internal, subjective effects can serve as discriminative stimuli and exert discriminative control over behavior once animals are trained. The discriminative stimulus effects of opioid analgesics, or mu-opioid receptor (MOR) agonists, have been well-characterized in rodents, monkeys, and humans (for example: rodent, Colpaert & Janssen, 1986; Pournaghash & Riley, 1993; Shannon & Holtzman, 1976; Walker & Young, 1993; monkey: DeRosset & Holtzman, 1986; Gerak & France, 1996; Negus, Picker, & Dykstra, 1991, human: Comer et al., 2008; 2009; 2010; Lasagna et al., 1955; Zacny, 2001), and all MOR agonists share similar discriminative stimulus properties. For example, in rats trained to discriminate an injection of morphine from saline, drugs with lower intrinsic efficacy (e.g., buprenorphine) partially generalize to the discriminative stimulus effects of morphine (Jones, Bigelow, & Preston, 1999; Brandt et al., 1997). However, compounds from other drug classes (e.g., pentobarbital) do not produce morphine-like responding (Broadbent et al., 1995; Colpaert, 1999). Importantly, drug discrimination assays can also be used to evaluate pharmacological phenomena, such as tolerance and dependence (Colpaert et al., 1976; Young, Kapitsopoulos, & Makhay, 1991; Paronis & Holtzman, 1985), and to dissect the overlapping mechanisms produced by compound pharmacological cues (e.g., Wolff & Leander, 1997).

The discriminative stimulus effects of opioid analgesics may also be altered by acute and chronic pain. For example, several studies in humans have demonstrated that exposure to escapable, noxious stimuli (e.g., cold pressor pain) decrease reported pleasant subjective effects of prescription opioid analgesics (Comer et al., 2010; Zacny & Bekman, 2004). In mice, acute administration of a chemical noxious stimulus (0.4% acetic acid, i.p.) produced a ∼2.2 fold rightward shift in the morphine dose effect curve in male, but not female, mice (Neelakantan, et al., 2015). These data collectively suggest exposure to a noxious stimulus may alter the interoceptive effects of MOR agonists. However, we know relatively little about whether or not chronic, ongoing/inescapable pain alters the discriminative stimuli of MOR agonists.

While opioid analgesics are not generally warranted for the treatment of chronic pain, data suggest that more than half (∼69%) of chronic neuropathic pain patients receive opioid analgesic treatment (Hoffman et al., 2017). Therefore, the goal of this study was to assess the extent to which a long-lasting pain state altered the discrimination of the MOR agonist fentanyl from saline or a non-opioid drug of abuse (cocaine) from saline in both male and female rats. In order to induce chronic pain-like states, we utilized the spared nerve injury (SNI) model developed by Decosterd & Wolfe (2000); this model was chosen for the persistent hyperalgesic-like state induced. Prior studies have demonstrated this state lasts at least 8 months following injury, providing ample time to evaluate potential pain-induced shifts in dose response curves for fentanyl (or cocaine) discrimination experiments (Erichsen & Blackburn-Munro, 2008).

## Methods

### Animals

Female and male Sprague-Dawley rats were purchased from Envigo laboratory (Indianapolis, IN). Following arrival, rats were group-housed and allowed at least one week of acclimation. All animals were housed in clear plastic cages with corncob bedding. Animal housing facilities were maintained on 12 hour light:dark cycle with lights on at 07:00. Seven to ten days after arrival, rats were single housed for 3-4 days with food and water available ad libitum. Rats were at least eight weeks old at the start of experiments. Approximately 72 hours prior to initiation of operant training, rats were single-housed, food restricted with males given ∼18 grams and females given ∼14 grams standard rat chow (5LOD, LabDiet, Tuscon, AZ) daily, and given 20 sucrose pellets (45 mg pellets, unflavored; F0023; Bioserv, Flemmington NJ) in their home cage.

### Surgery

The spared nerve injury (SNI) or sham surgeries were performed as described by Decosterd & Wolfe (2000). Briefly, rats were anesthetized with 90 mg/kg ketamine (i.p.) and 10 mg/kg xylazine (i.p.). Carprofen (5 mg/kg, s.c.) was given prior to surgery as pre-operative analgesia as well as 24- and 48 hr after surgery for post-operative analgesia. An incision was made in the left femoral muscle to expose the sciatic nerve. In the SNI condition, a 2 mm portion of the tibial and peroneal branches was removed while the sural branch was left unmanipulated. In the sham condition, the femoral muscle incision was made, but there was no manipulation of any branch of the sciatic nerve. In both surgeries, the muscle and skin incision were closed with absorbable suture and surgical staples, respectively.

### Apparatus

All experiments were conducted in 18 standard Med PC operant chambers (ENV-008CT, Med Associates, St. Albans, VT) housed inside ventilated sound-attenuating chambers (ENV-018CT). Chambers were equipped with two illuminated nosepoke devices (ENV-114BM), a food hopper, a pellet dispenser, a white house light, and three panel stimulus lights. Nosepoke manipulandum were located on the right side of the chamber on either side of the pellet hopper (ENV-200R2M). Three-panel LED stimulus lights were located above both nosepokes (ENV-114BM). The pellet hopper was connected to a pellet dispenser (ENV-203-45) filled with 45 mg dustless, unflavored sucrose pellets (F0023, Bioserv, Flemmington, NJ). A white house light was located on the left side of the chamber.

### Procedure

#### Operant training

Training, testing, and maintenance sessions were modeled on previous work (Jutkiewicz et al., 2011). Illumination of the lights in the nosepokes signaled the initiation of response periods. First, animals were trained to respond for 45 mg sucrose pellets in a 20-min session in which responding on either nosepoke resulted in sucrose pellet delivery and illumination of the house light. The response requirement was gradually increased from fixed ratio (FR)1 to FR10 over time. Next, a 5-min blackout period was introduced at the beginning of the session, which increased to 15 min over 6-8 sessions. Simultaneously, the responding period decreased from 20 min to 5 min over 5-10 sessions. Therefore, a single component was comprised of a 15 min blackout and 5 min response (S^D^) period. At this time, individual preferences for left or right nosepoke manipulanda were determined.

#### Discrimination Training

Training drug dose was assigned to the preferred nosepoke in ∼50% of rats. Rats were trained to discriminate 0.032 mg/kg fentanyl (s.c.) from saline or 5.6 mg/kg cocaine (i.p.) from saline. Saline or training drugs were administered by the experimenter immediately prior to the start of each component. Following 15 min of blackout, responding on a FR10 schedule of reinforcement on the injection-appropriate nosepoke was reinforced with 45 mg sucrose pellet delivery during the 5-min S^D^ period o. Each pellet delivery was followed by a 10 sec timeout. Completing an FR on the incorrect nosepoke resulted in a 10 sec timeout with no pellet delivery. At the end of the 5-min S^D^ period, the rat was removed from the operant chamber and injected with either saline or drug and returned immediately to the chamber for the subsequent component. The number of components was randomized across days, ranging from 1 – 4 components. Multiple component training consisted of 1 – 4 injections of saline (saline, saline-saline, saline-saline-saline, or saline-saline-saline-saline) or 0-3 injections of saline followed by administration of training dose (drug, saline-drug, saline-saline-drug, saline-saline-saline-drug). Training dose was always administered in the last component.

Discrimination training continued until the following criteria were met: 1) >80% injection appropriate responding across components and 2) successful completion of first FR on injection-appropriate nosepoke. Testing sessions (as described below) were not initiated until training criteria were met for 5 consecutive days.

#### Testing & Maintenance

Test sessions were conducted no more than 2 times per week; however, if training criteria were not met on any maintenance day, then three subsequent maintenance days were required in which the aforementioned criteria were met prior to re-initiating testing. Test sessions consisted of four components unless an animal failed to complete a single FR in components 1-3. Drugs were administered immediately prior to the start of each component as described above, and dose response curves were constructed by administering cumulative doses of drug at the start of each component. Full or complete generalization to a discriminative cue was defined as >85% of responding on the drug-associated nose poke and completing at least one FR.

#### Experimental Design

In all rats, operant and discrimination (fentanyl vs saline *or* cocaine vs saline) training and dose effect curves for training drug and substitutions were evaluated repeatedly prior to surgery. Once initial dose effect curves were determined, SNI or sham surgery was performed. Rats were given 72 hours post-operative recovery as described above. Then, dose effect curves for training drug and substitutions were repeatedly evaluated over 4 mo.

#### Drugs

Cocaine and ketamine were obtained from the Univ Michigan Medical Hospital Pharmacy. Fentanyl, nalbuphine, and gabapentin were purchased from Sigma Aldrich (St. Louis, MO). Morphine was purchased as a 50 mg/mL solution from Henry Schein. SNC80 was generously gifted by Dr. Kenner C. Rice. Naltrexone, and quinpirole were obtained from Tocris Biosciences (Bristol, UK). Xylazine was obtained from Heartland Vet Supply (Hastings, NE). Carprofen (Rimadyl) was purchased from Zoetis (Parsippany, NJ).

Fentanyl citrate, morphine sulphate, cocaine hydrochloride, nalbuphine hydrochloride, gabapentin, naltrexone hydrochloride, quinpirole hydrochloride, and amphetamine sulfate were dissolved in sterile, physiological saline. Carprofen and xylazine were diluted in sterile water. SNC80 was dissolved in 3-5% 1 M HCl. SNC80 vehicle was 3% 1M HCl, and vehicle for all other drugs tested in discrimination procedures was saline.

Amphetamine, cocaine, gabapentin, ketamine, nalbuphine, quinpirole, and xylazine were given intraperitonially (i.p.) and all other drugs were delivered subcutaneously (s.c.).

#### Statistical Analysis & Data Presentation

All experiments were conducted according to a pre-determined experimental design, and there are 6-8 data points in every statistical group. Dose effect curves are presented as an average for each testing period: pre-surgery, 1-2, and 3-4 months post-surgery. Statistics were conducted in SPSS version 29. Data were analyzed using 4-way repeated measure ANOVAs with the following variables: sex (male, female); surgical status (sham, SNI); dose (dosages varied across drugs but vehicle, 3-4 doses); and time (pre surgery, 1-2 or 3-4 months after surgery). Necessary corrections were made according to established Mauchly’s W criteria. Analyses were conducted on % training drug responding as well as rates of responding for each drug, over time in cocaine or fentanyl trained groups. Main effects and interaction terms necessary for interpretation are reported. To further examine main effect of time or dose, post hoc one-way ANOVAs with post hoc analyses were conducted. In these analyses, data were collapsed across non-significant main effects when possible (e.g., non-significant main effect of sex, data collapsed across sex).

## Results

Figure 1. % Fentanyl Responding

Fentanyl produced dose-dependent interoceptive effects in both males (1A, D) and females (1F, J) prior to and after surgery, supported by a main effect of dose [F(3, 60)=39.81, p<0.001, 17^2^_p_ =0.67]. There was no main effect of time [F(2, 40)=27.32, p=0.1, 17^2^_p_ =0.39], sex [F(1, 20)=0.01, p=0.92, 17^2^_p_ =0.001], or surgical status [F(1, 20)=2.69, p=0.12, 17^2^_p_ =0.09]. However, there was a three-way interaction between time, dose, and surgical status [F(6, 120)=2.93, p=0.01, 17^2^_p_ =0.13]. This is likely explained by the small rightward shifts in the dose effect curves following surgery, and these are slightly greater in injured rats (Fig. 1D, J) than sham rats (Fig. 1A, F), particularly in male rats (Fig 1A, D). All other interaction terms failed to reach significance (p’s>0.07). Collectively, these data suggest small, rightward shifts in the fentanyl dose effect curves in both sham and SNI rats.

**Figure 1.**
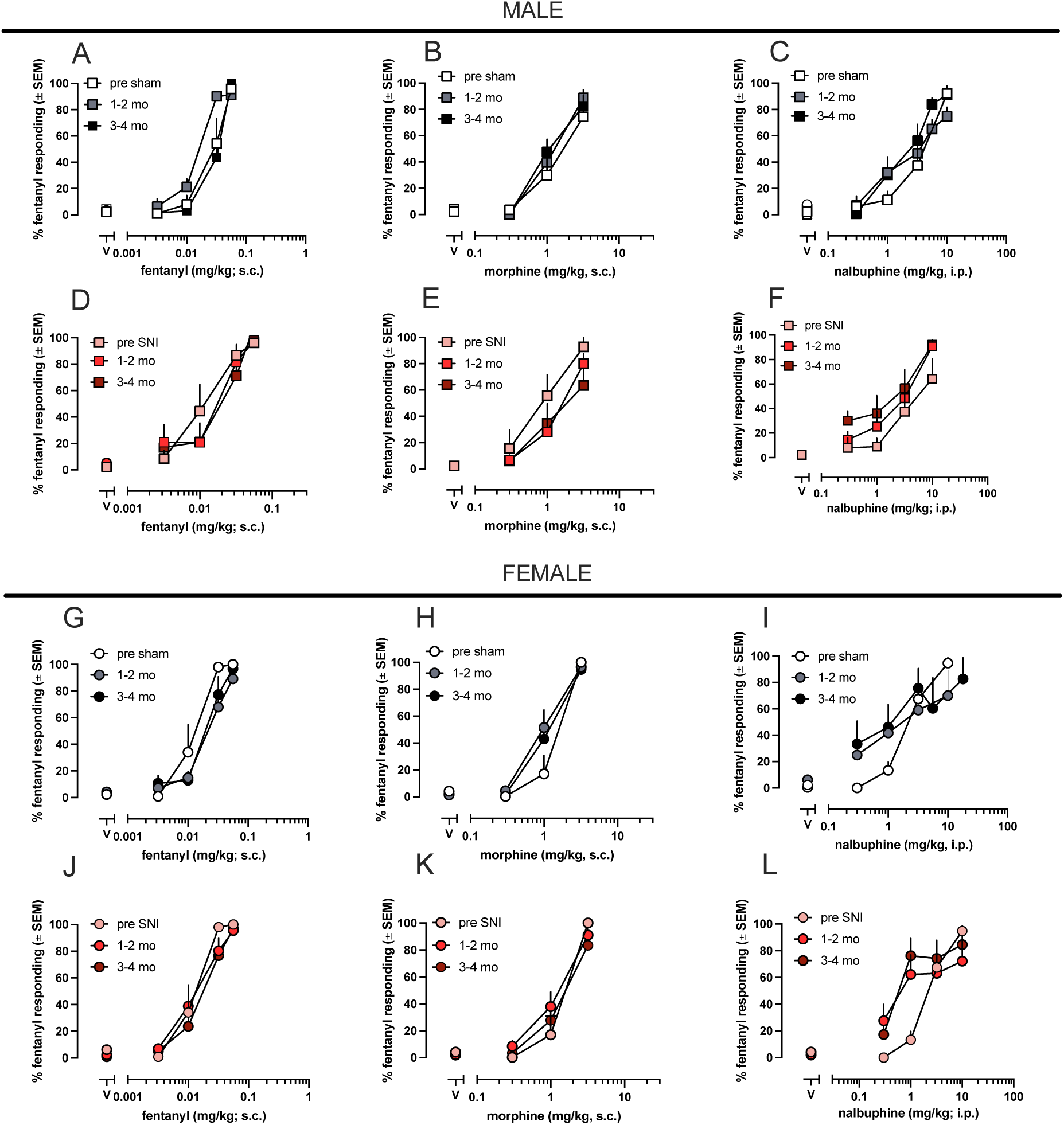
Interoceptive effects of opioid analgesics in males and females in the presence or absence of chronic neuropathic pain. In males, prior to either sham (A) or SNI (D) surgery fentanyl produced dose dependent increases in % fentanyl responding, and there were no significant shifts in the dose effect curves after up to 4 months of sham (A) or SNI surgery (D). Morphine produced dose dependent increases in % fentanyl responding in male rats prior to sham (B) or SNI (E) surgery, and there were no significant shifts in the dose effect curves following sham (B) or SNI (E) surgery. Nalbuphine produced dose dependent increases in % fentanyl responding prior to sham (C) or SNI (F) surgery, and in both sham (C) or SNI (F) groups, leftward shifts were observed in the dose effect curves following surgery. In females, prior to either sham (G) or SNI (J) surgery fentanyl produced dose dependent increases in % fentanyl responding, and there were no significant shifts in the dose effect curves after up to 4 months of sham (G) or SNI surgery (J). Morphine produced dose dependent increases in % fentanyl responding in male rats prior to sham (H) or SNI (K) surgery, and there were no significant shifts in the dose effect curves following sham (H) or SNI (K) surgery. Nalbuphine produced dose dependent increases in % fentanyl responding prior to sham (H) or SNI (L) surgery, and in both sham (H) or SNI (L) groups leftward shifts were observed in the dose effect curves following surgery.

Morphine dose-dependently substituted for fentanyl in both males (Fig. 1B, E) and females (Fig. 1H, K) prior to and after surgery, supported by a main effect of dose [F(2.16, 38.82)=345.64, p<0.001, 17^2^_p_ =0.95]. There was no main effect of time [F(2, 36)=0.17, p=0.84, 17^2^_p_ =0.009],surgical status [F(1, 18)=0.39, p=0.54, 17^2^_p_ =0.02], or sex [F(1, 18)=2.93, p=0.1, 17^2^_p_ =0.14]. Though, there was a two-way interaction between dose and surgical status [F(2.16, 38.82)=3.38, p=0.04, 17^2^_p_ =0.16] such that there were small, rightward shifts in the dose effect curves in injured animals compared sham rats. All other interaction terms failed to reach significance (p’s>0.07). Collectively, these results suggest that there were small rightward shifts in the morphine dose effect curves in male injured animals.

Nalbuphine dose dependently substituted for fentanyl in both male (Fig. 1C, F) and female (Fig. 1G, J) rats prior to and after surgery [F(2.45, 46.52)=27.02, p<0.001, 17^2^_p_ =0.59]. There was no main effect of sex [F(1, 19)=2.73, p=0.12, 17^2^_p_ =0.13], surgical status [F(1, 19)=0.89, p=0.36, 17^2^_p_ =0.045], or time [F(2, 38)=1.38, p=0.27, 17^2^_p_ =0.07]. However, there was a three-way interaction between dose, time, and sex [F(8, 152)=1.78, p=0.08, 17^2^_p_ =0.09], which is likely explained by the observed increases in sensitivity (i.e., leftward shift in the dose effect curve) to the discriminative stimulus effects of nalbuphine. Though, there was no dose by time by surgical status three-way interaction, suggesting the increased sensitivity to nalbuphine substitution is unrelated to SNI-induced hypersensitivity. The smallest increase in sensitivity was observed in the male sham group, but this effect was not significant [F(8, 152)=0.25, p=0.98, 17^2^_p_ =0.013]. All other interaction terms failed to reach significance (p’s> 0.12). Collectively, these results suggest a change in sensitivity to nalbuphine over time independent of nerve injury status.

Figure 2. Rates of Responding in Fentanyl-Trained Animals

Fentanyl dose dependently decreased rates of responding prior to and after surgery in both males (Fig. 2A, D) and females (Fig. 2G, J), reflected by a main effect of dose [F(2.45, 47.00)=56.83, p<0.001, 17^2^_p_ =0.75]. There was no main effect of time [F(1.49, 28.25)=0.03, p=0.93, 17^2^_p_ =0.002], surgical status [F(1, 19)=1.83, p=0.19, 17^2^_p_ =0.09], or sex [F(1, 19) = 0.001, p=0.97, 17^2^_p_ =0]. However, there was a significant dose by sex interaction [F(2.47, 47.00)=5.87, p=0.003, 17^2^_p_ =0.24], suggesting the rate decreasing effects are slightly more potent in female rats. There was no three-way interaction between sex, dose, and surgical status [F(2.47, 47.00)=0.57, p=0.61, 17^2^_p_ =0.03], suggesting that SNI-induced hypersensitivity is not responsible for this small sex difference. All other interaction terms failed to reach significance (p’s>0.16). These data collectively suggest SNI-induced hypersensitivity did not induce a change in fentanyl-induced rate suppressant effects.

**Figure 2.**
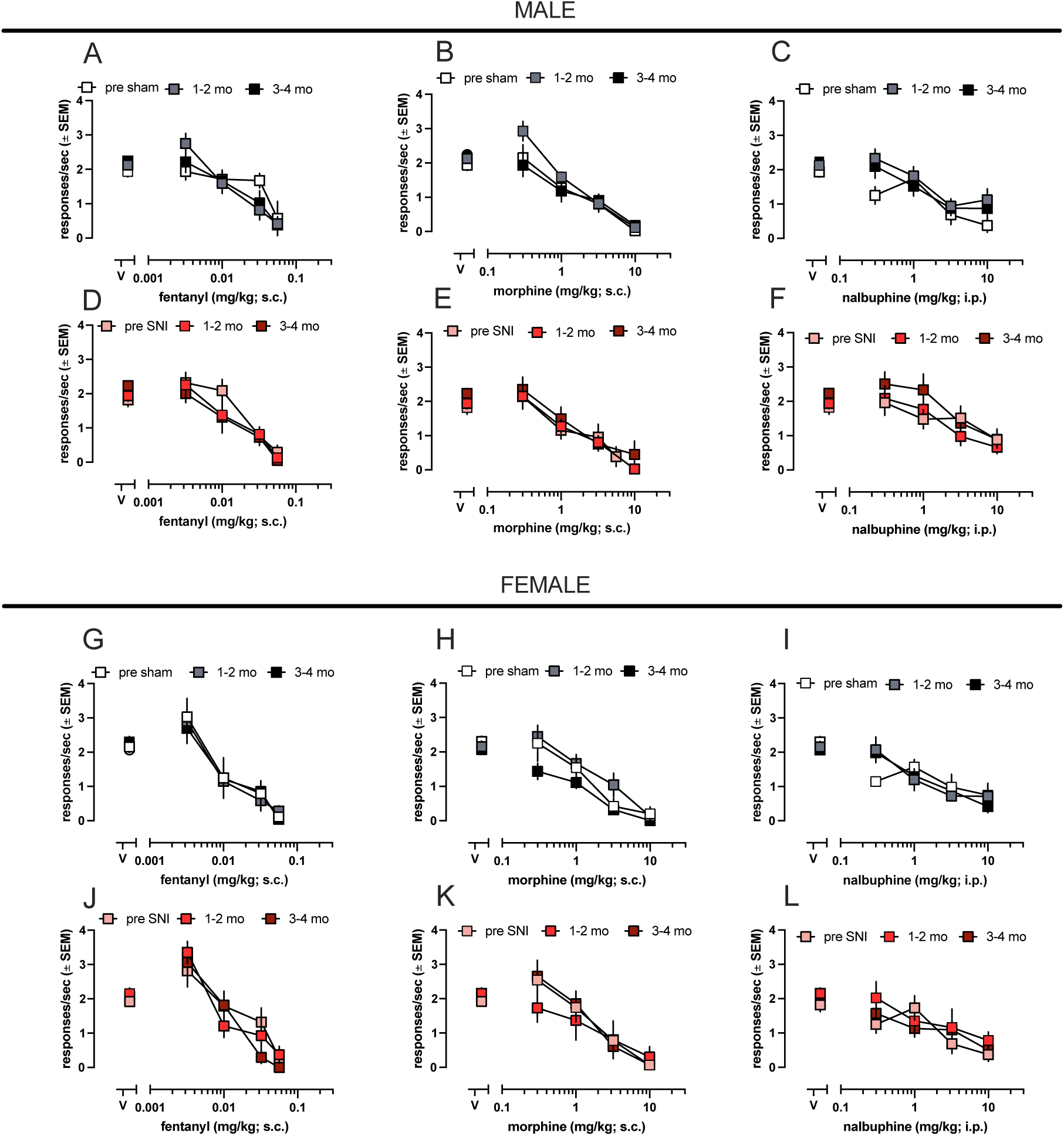
Rate suppressant effects of opioid analgesics in the presence or absence of chronic neuropathic pain. In males, prior to either sham (A) or SNI (D) surgery fentanyl produced dose dependent decreases in rates of responding, and there were no significant shifts in the dose effect curves after up to 4 months of sham (A) or SNI surgery (D). Morphine produced dose dependent decreases in rates of responding in male rats prior to sham (B) or SNI (E) surgery, and there were no significant shifts in the dose effect curves following sham (B) or SNI (E) surgery. Nalbuphine produced dose dependent decreases in rates of responding prior to sham (C) or SNI (F) surgery, and in both sham (C) or SNI (F) groups, leftward shifts were observed in the dose effect curves following surgery. In females, prior to either sham (G) or SNI (J) surgery fentanyl produced dose dependent decreases in rates of responding, and there were no significant shifts in the dose effect curves after up to 4 months of sham (G) or SNI surgery (J). Morphine produced dose dependent decreases in rates of responding in male rats prior to sham (H) or SNI (K) surgery, and there were no significant shifts in the dose effect curves following sham (H) or SNI (K) surgery. Nalbuphine produced dose dependent decreases in rates of responding prior to sham (H) or SNI (L) surgery, and in both sham (H) or SNI (L) groups leftward shifts were observed in the dose effect curves following surgery.

Morphine dose dependently decreased rates of responding prior to and after surgery in both males (Fig. 2B, E) and females (Fig. 2H, K), reflected by a main effect of dose [F(2.29, 45.86)=51.88, p<0.001, 17^2^_p_ =0.72]. There was no main effect of time [F(2, 40)=0.70, p=0.50, 17^2^_p_ =0.03], sex [F(1, 20)=0.33, p=0.57, 17^2^_p_ =0.02], or surgical status [F(1, 20=0.03, p=0.87, 17^2^_p_ =0.001], suggesting morphine induced rate suppressant effects did not largely differ across conditions and that SNI-induced hypersensitivity did not alter morphine-induced rate suppressant effects. All interaction terms failed to reach significance (p’s>0.11).

Nalbuphine dose-dependently decreased rates of responding prior to and after surgery in both males (Fig. 2C, F) and females (Fig. 2I, L), reflected by a main effect of dose [F(2.31, 43.93)=27.05, p<0.001, 17^2^_p_ =0.59]. There was no main effect of time [F(2, 38)=0.82, p=0.45, 17^2^_p_ =0.04], sex [F(1, 19)=2.41, p=0.14, 17^2^_p_ =0.11], or surgical status [F(1, 19)=1.02, p=0.32, 17^2^_p_ =0.05]. However, there was a significant interaction between time and dose, suggesting that some doses of nalbuphine were less rate suppressant over time (Fig. 2C, F, I, L). There was no three-way interaction between time, dose, and surgical status [F(4.33, 82.35)=0.39, p=0.82, 17^2^_p_ =0.02], indicating that the small shifts in nalbuphine dose response curves over time were not related to nerve injury. All other interaction terms failed to reach significance (p’s>0.19).

Figure 3. Cocaine-induced interoceptive effects in presence or absence of chronic neuropathic pain

Cocaine produced dose-dependent interoceptive effects in both males (Fig. 3A, C) and females (Fig. 3G, K), reflected by a main effect of dose [F(4, 68)=120.59, p<0.001, 17^2^_p_ =0.88]. There was no main effect of time [F(2, 34)=1.08, p=0.35, 17^2^_p_ =0.06], sex [F(1, 22)=0.54, p=0.86, 17^2^_p_ =0.002], or surgical status [F(1, 22)= 1.53, p=0.48, 17^2^_p_ =0.03], suggesting cocaine-induced interoceptive effects were not altered over time, by sex, or by SNI-induced hypersensitivity. However, there is a three-way time by dose by sex interaction [F(4.00, 68.06)=0.89, p=0.06, 17^2^_p_ =0.13], suggesting that cocaine was more potent in male than female rats prior to surgery. All other interaction terms failed to reach significance p’s>0.13.

**Figure 3.**
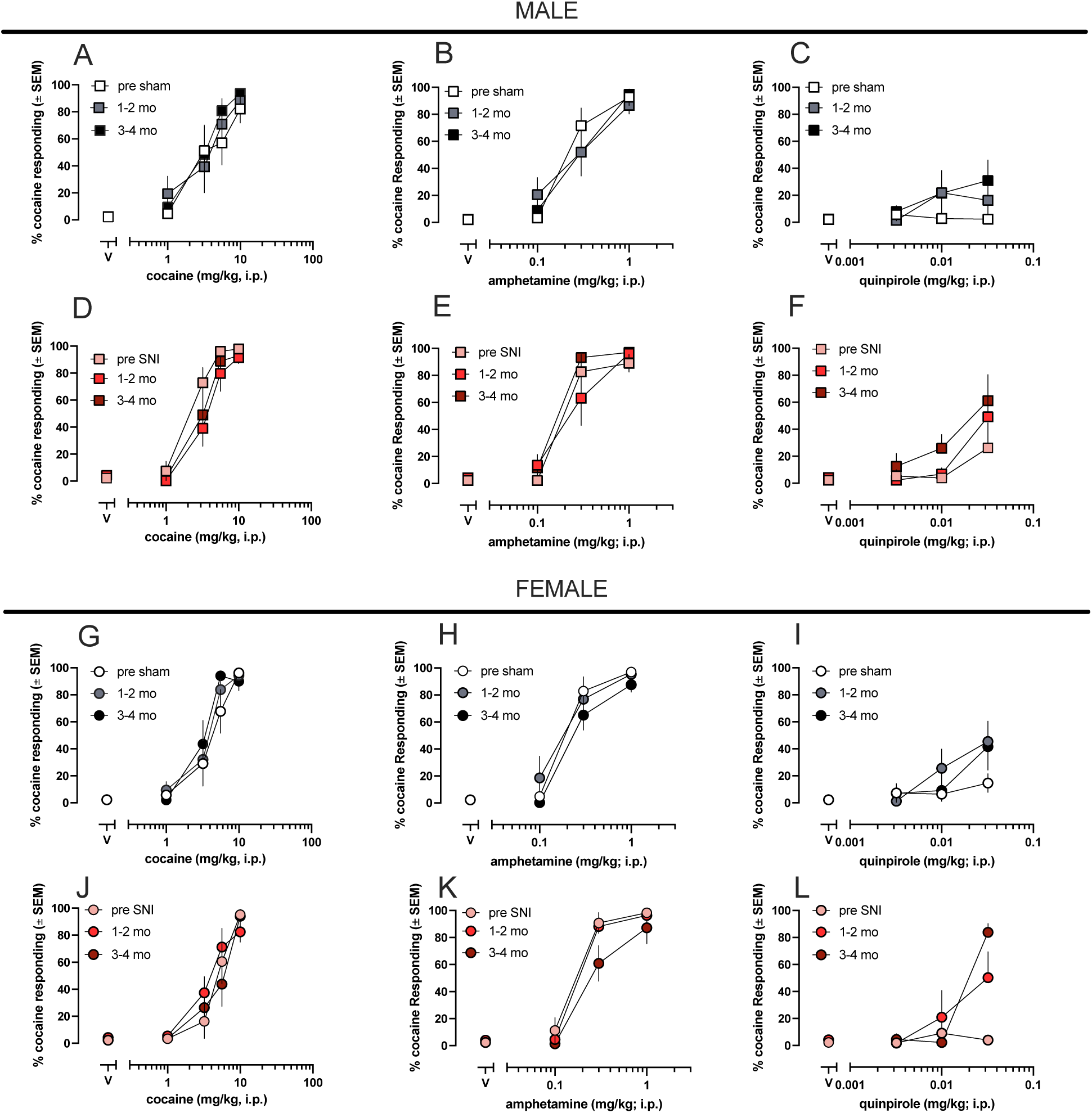
Interoceptive effects of dopaminergic drugs in the presence or absence of chronic neuropathic pain. In males, prior to either sham (A) or SNI (D) surgery cocaine produced dose dependent increases in % cocaine responding, and there were no significant shifts in the dose effect curves after up to 4 months of sham (A) or SNI surgery (D). Amphetamine produced dose dependent increases in % cocaine responding in male rats prior to sham (B) or SNI (E) surgery, and there were no significant shifts in the dose effect curves following sham (B) or SNI (E) surgery. Prior to sham (C) or SNI (F) surgery, quinpirole produced low levels (<25%) of % cocaine responding. Following sham (C) or SNI (F) surgery, quinpirole produced significantly higher levels of % cocaine responding in sham (C) and SNI (F) groups, albeit lower levels in sham. In females, prior to either sham (G) or SNI (J) surgery, cocaine produced dose dependent increases in % cocaine responding, and there were no significant shifts in the dose effect curves after up to 4 months of sham (G) or SNI surgery (J). Amphetamine produced dose dependent increases in % cocaine responding in female rats prior to sham (H) or SNI (K) surgery, and there were no significant shifts in the dose effect curves following sham (H) or SNI (K) surgery. Prior to sham (I) or SNI (L) surgery, quinpirole produced low levels (<25%) of % cocaine responding. Following sham (I) or SNI (L) surgery, quinpirole produced significantly higher levels of % cocaine responding in sham (I) and SNI (L) groups, albeit lower levels in sham.

Amphetamine dose-dependently substituted for cocaine in both males (Fig. 3B, E) and females (Fig. 3H, K), reflected by a main effect of dose [F(2.20, 26.38)=17.80, p<0.001, 17^2^_p_ =0.6]. There was no main effect of time [F(2, 24)=1.76, p0.19, 17^2^_p_ =0.13], sex [F(1, 22) = 1.76, p=0.21, 17^2^_p_ =0.13], or surgical status [F(1, 22)=0.26, p=0.62, 17^2^_p_ =0.02], suggesting that discriminative stimulus effects of amphetamine was not altered over time, by sex, or by SNI-induced hypersensitivity. Though, there was a two-way dose by time interaction, likely reflecting the small rightward shift in the amphetamine dose effect curves at some time point over the study. There was also a significant two-way dose by sex interaction [F(2.20, 26.38)=3.79, p=0.03, 17^2^_p_ =0.24], likely reflecting the slightly higher potency in female rats. All other interaction terms failed to reach significance p’s>0.13.

Quinpirole substitution produced low levels of cocaine appropriate responding prior to surgery in both males (Fig. 3C, F) and females (Fig. 3I, L). After surgery, quinpirole substitution produced higher levels of cocaine appropriate responding in both sham and SNI groups, though to a greater extent in SNI groups. Quinpirole substitution was dose dependent, reflected by a main effect of dose [F(2.17, 22.86)=8.71, p= 0.001, 17^2^_p_ =0.44]. There was no main effect of sex [F(1, 11) = 0.18 p0.68, 17^2^_p_ =0.016] or surgical status [F(1, 11)=0.995, p=0.34, 17^2^_p_ =0.08]. There was no main effect of time [F(1.51, 16.63)=1.08, 0.34, 17^2^_p_ =0.09], though there was a significant interaction between dose and time [F(6, 66)=15.23, p<0.001, 17^2^_p_ =0.58], l and significant three-way interaction between dose, time, and surgery status [F(6, 66)=3.40, p=0.006, 17^2^_p_ =0.24], reflecting greater increases in quinpirole substitution for cocaine over time in the injured animals than sham (male Fig. 3C, F; female Fig. 3I, L).

Figure 4. Rates of responding in cocaine-trained animals

Cocaine dose-dependently decreased rates of responding prior to and after surgery in both male (Fig. 4A, D) and female rats (Fig. 4G, K), reflected by a main effect of dose [F(2.70, 54.06)=77.81, p<0.001, 17^2^_p_ =0.80]. There was no main effect of time [F(2, 40)=1.63, p=0.21, 17^2^_p_ =0.8] or surgical status [F(1, 20)=0.73, p=0.40, 17^2^_p_ =0.04]. Cocaine more potently suppressed rates of responding in female rats than male rats, reflected by a main effect of sex [F(1, 20)=8.56, p=0.008, 17^2^_p_ =0.3]. All interaction terms failed to reach significance (p’s>0.18). These data collectively indicate that SNI-induced hypersensitivity did not alter cocaine-induced decreases in rates of responding.

**Figure 4.**
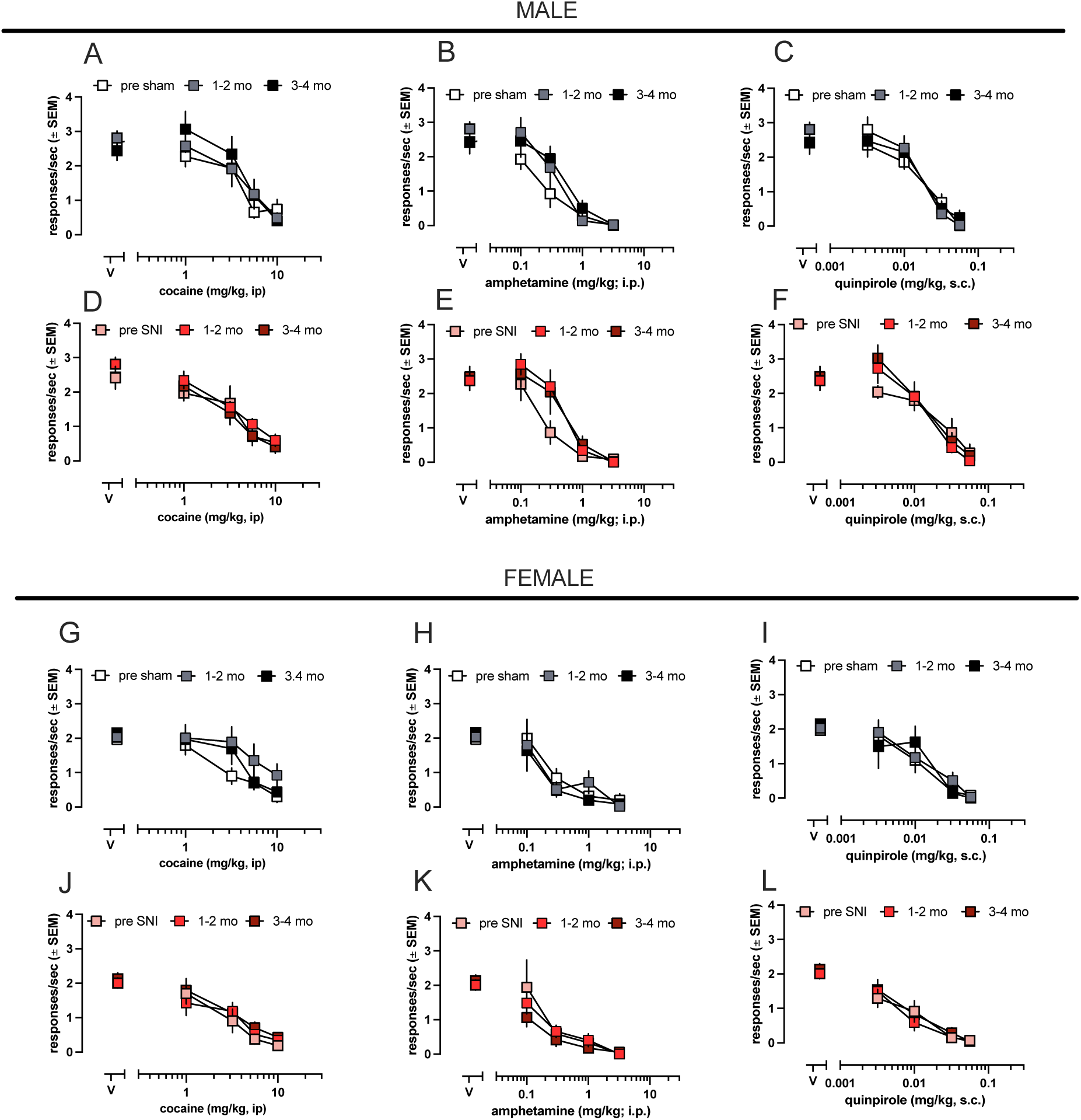
Rate suppressant effects of dopaminergic drugs in the presence or absence of chronic neuropathic pain. In males, prior to either sham (A) or SNI (D) surgery cocaine produced dose dependent decreases in rates of responding, and there were no significant shifts in the dose effect curves after up to 4 months of sham (A) or SNI surgery (D). Amphetamine produced dose dependent decreases in rates of responding in male rats prior to sham (B) or SNI (E) surgery, and there were no significant shifts in the dose effect curves following sham (B) or SNI (E) surgery. Quinpirole produced dose dependent decreases in rates of responding prior to sham (C) or SNI (F) surgery, and in both sham (C) or SNI (F) groups, no significant shifts were observed in the dose effect curves following surgery. In females, prior to either sham (G) or SNI (J) surgery cocaine produced dose dependent decreases in rates of responding, and there were no significant shifts in the dose effect curves after up to 4 months of sham (G) or SNI surgery (J). Amphetamine produced dose dependent decreases in rates of responding in male rats prior to sham (H) or SNI (K) surgery, and there were no significant shifts in the dose effect curves following sham (H) or SNI (K) surgery. Quinpirole produced dose dependent decreases in rates of responding prior to sham (H) or SNI (L) surgery, and in both sham (H) or SNI (L) groups no significant shifts were observed in the dose effect curves following surgery.

Amphetamine dose-dependently decreased rates of responding prior to and after surgery in both male (Fig. 4B, D) and female rat (Fig. 4H, K), reflected by a main effect of dose [F(2.28, 45.51)=71.92, p<0.001, 17^2^_p_ =0.78]. There was no main effect of time [F(2, 40)=1.62, p=0.21, 17^2^_p_ =0.08], surgical status [F(1, 20)=0.02, p=0.88, 17^2^_p_ =0.001], or sex [F(1, 20)=23.70, p=0.054, 17^2^_p_ =0.50]. The main effect of sex was close to significant, and there was a two-way interaction between dose and sex [F(2.28, 45.51)=6.91, p=0.002, 17^2^_p_ =0.26], such that amphetamine was more rate suppressant in females (Fig. 4H, K) than males (Fig. 4B, E). All other interaction terms failed to reach significance (p’s>0.27). Collectively these data suggest chronic neuropathic pain did not alter amphetamine-induced rate suppressant effects.

Quinpirole dose-dependently decreased rates of responding prior to and after surgery in both male (Fig. 4C, F) and female rats (Fig. 4I, L), reflected by a main effect of dose [F(2.03, 40.61)=77.44, p<0.001, 17^2^_p_ =0.80]. There was no main effect of time [F(2, 40)=0.16, p=0.85, 17^2^_p_ =0.008] or surgical status [F(1, 20)=0.86, p=0.36, 17^2^_p_ =0.04]. There was a main effect of sex [F(1, 20)=24.21, p<0.001, 17^2^_p_ =0.55] and a significant [two-way interaction between dose & sex: F(2.03, 40.61)=5.62, p=0.007, 17^2^_p_ =0.22], such that female rats (Fig. 4I, L) were more sensitive than males (Fig. 4C, F) to the rate decreasing effects of quinpirole. All other interaction terms failed to reach significance (p’s>0.21). Collectively these results suggest chronic neuropathic pain did not alter quinpirole-induced rate suppressant effects.

## Discussion

The goal of the present study was to determine if chronic neuropathic pain altered the discriminative stimulus of opioid analgesics. Several previous studies in humans reported that acute, laboratory-induced pain states decreased the magnitude of the pleasant subjective effects of opioids (Comer et al., 2010; Zacny & Bekman, 2004); however, we do not fully understand how chronic pain states may alter opioid-induced pain states. Discrimination training and pre-surgical testing of substitution dose-effect curves were completed prior to and after induction of chronic neuropathic pain or sham states over 4 months. We repeatedly evaluated dose effect curves to determine if, within-subject, chronic pain states alter the interoceptive effects of opioid analgesics over time.

First, as expected, fentanyl produced dose-dependent interoceptive effects in both male and female rats (Fig. 1) and dose-dependently decreased rates of responding (Fig. 2). Following sham or SNI surgeries, there were minor rightward shifts in the fentanyl interoceptive effects dose response curves, suggesting a small, but significant decrease in fentanyl-induced interoceptive effects independent of surgical status. There was no significant change in fentanyl-induced rate suppressant effects in any groups over time (Fig. 2). Prior to and after surgery, morphine dose-dependently substituted for fentanyl in both males and females, as expected (Fig. 2). Similar to what was observed with fentanyl, there were small but significant changes in effectiveness of individual doses of morphine 3-4 months after sham *or* SNI surgery in male rats, suggesting SNI-induced hypersensitivity failed to alter the substitution of morphine to the fentanyl discriminative stimulus. Collectively, these data suggest that SNI-induced hypersensitivity failed to alter the discriminative stimulus of fentanyl or morphine (high efficacy MOR agonists) in both male and female rats. These changes may are unlikely to be explained by previously reported pain-induced decreased expression and activity of MORs (Back et al., 2006; Campos-Jurado et al, 2019; Dong et al., 2019; Hou et al., 2017; Ji et al., 1995; Kaneuchi et al., 2019; Pol et al., 2006; Porecca et al., 1998; Thompson et al., 2018; Yamamoto et al., 2008; Zhang et al., 1998), as small shifts were observed in the sham groups as well (Fig. 1). These findings are inconsistent with previous human studies that have demonstrated that acute pain states decreased the magnitude of opioid analgesic-induced subjective effects (Comer et al., 2010; Zacny & Beckham, 2004); however, methodological differences such as type of pain (human-acute, escapable; present study-chronic, inescapable) may explain the observed differences in results.

Nalbuphine, a partial MOR agonist, dose-dependently substituted for fentanyl prior to and after surgery in both males and females (Fig. 1) and dose-dependently decreased rates of responding (Fig. 2) as expected. After surgery, in both sham and SNI groups, nalbuphine was more potent, suggesting an increase in sensitivity to nalbuphine-induced interoceptive effects over time that is independent of SNI-induced hypersensitivity. Therefore, other factors may be responsible for the increased sensitivity to the discriminative stimulus of nalbuphine over time.

Rats began these studies in age-matched cohorts, but these studies were longitudinal, and animals ended the experiments significantly older (∼12 months) than they started (∼2 months), so it is possible that increased age of subjects may contribute to the changes in sensitivity to the nalbuphine discriminative stimulus over time. Increased sensitivity to MOR agonist-induced subjective effects has been observed in older humans compared to younger humans (Cherrier et al., 2009; McLachlan et al., 2011; Scott et al., 1987), suggesting age increases sensitivity to opioid-analgesic induced subjective effects. The leftward shift in the nalbuphine dose effect curve is potentially related to increased age of animals over the course of discrimination experiments. this increased sensitivity to the discriminative stimulus of nalbuphine is consistent with a previous study showing increased effectiveness of nalbuphine in 24-month old rats compared to 2-month old rats in the warm water tail withdrawal assay, suggesting there is an increased sensitivity to nalbuphine-induced antinociceptive-like effects with increased age (Smith & Grey, 2001). It is possible that we only observed an increased sensitivity in the discriminative stimulus of nalbuphine (but not morphine, fentanyl) as these assays may not be sensitive enough to detect small changes in higher efficacy MOR agonists. Future studies should carefully evaluate the reinforcing effects of partial MOR agonists over time in the presence or absence of pain states.

Collectively, SNI-induced hypersensitivity failed to alter the discriminative stimuli of MOR agonists (fentanyl, morphine, nalbuphine) (Fig. 1). While evaluating the impact of chronic neuropathic pain states on the interoceptive effects of opioid analgesics was the primary goal of this study, we also examined the interoceptive effects of cocaine, an indirect dopaminergic agonist and a non-opioid drug of abuse. In animals trained to discriminate 5.6 mg/kg cocaine from saline, cocaine and amphetamine produced dose-dependent increases in cocaine-appropriate responding prior to and after sham or SNI surgery (Fig. 3) and dose-dependently decreased rates of responding (Fig. 4). No effects of SNI-induced hypersensitivity were observed on the interoceptive or rate decreasing effects of cocaine or amphetamine. Collectively, these results suggest that the interoceptive effects of indirect dopaminergic agonists were not altered by chronic neuropathic pain, suggesting no change in abuse potential. This is supported by the finding that chronic neuropathic pain did not alter the reinforcing effects of cocaine.

Prior to surgery, quinpirole produced low levels of cocaine-appropriate responding, consistent with previous studies (Katz & Witkin, 1982; Collins et al., 2014). After surgery, increased sensitivity to quinpirole was observed in all groups such that larger doses of quinpirole produced higher levels of cocaine-appropriate responding. While increased sensitivity was observed in sham *and* SNI groups, there was a greater increase in the SNI groups (Fig. 3); however, no changes in quinpirole-induced rate decreasing effects were observed in any group (Fig. 4). The increased sensitivity to the quinpirole discriminative stimulus in chronic neuropathic pain was somewhat surprising. Increased quinpirole-induced cocaine-appropriate responding in both sham and SNI groups suggests that while SNI-induced hypersensitivity may contribute, other factors likely are involved. Future studies should directly evaluate the reinforcing effects of dopaminergic agonists in the presence or absence of chronic neuropathic pain.

Overall, the results presented in this study suggest that chronic neuropathic pain failed to alter the discriminative stimuli of MOR agonists, though changes in sensitivity were observed over time and were independent of surgical status. The present study also found that the discriminative stimulus of cocaine or amphetamine was not altered by presence of SNI-induced hypersensitivity, though quinpirole was more cocaine-like in both sham and SNI groups over time. Therefore, in the future, it will be important to directly evaluate 1) the abuse potential of multiple partial MOR agonists in the presence or absence of chronic neuropathic pain, as well as in aged rats and 2) directly evaluate the abuse potential of dopaminergic agonists in the presence or absence of chronic neuropathic pain.

## Author Contributions

Participated in research design: Burgess, Traynor, Jutkiewicz

Conducted experiments: Burgess

Performed data analysis: Burgess, Jutkiewicz

Wrote or contributed to the writing of the manuscript: Burgess, Traynor, Jutkiewicz

## Footnotes

This work was supported by the Dr. Ben & Diana Lucchesi Graduate Education Fellowship (GEB); and the National Institutes of Health National Institute on Drug Abuse [Grants T32 DA007281 (GEB), UG3 DA056884 (JRT)].

